# Single-molecule analysis reveals the rapid effect of estradiol on the surface movement of AMPAR in live neurons

**DOI:** 10.1101/602813

**Authors:** Soma Godó, Klaudia Barabás, Ferenc Lengyel, Dávid Ernszt, Tamás Kovács, Miklós Kecskés, Csaba Varga, Tibor Z. Jánosi, Gergely Kovács, Barbara Orsolits, Takahiro Fujiwara, Akihiro Kusumi, István M. Ábrahám

## Abstract

The gonadal steroid 17β-estradiol (E2) rapidly alters glutamatergic neurotransmission, but its direct effect on the AMPA receptor (AMPAR) remains unknown. Live-cell single-molecule imaging experiments revealed that E2 rapidly and dose-dependently alters the surface movement of AMPAR via membrane estrogen receptors with distinct effects on somas and neurites. The effect of E2 on the surface mobility of AMPAR depends on the integrity of the cortical actin network.

## Main text

The gonadal steroid 17β-estradiol (E2) plays a role in a wide range of biological actions from fertility to neuroprotection (1,2). The cellular effects of E2 have been proposed to be mediated by slow transcriptional action through classic nuclear receptors (ERα and ERβ). In addition to its classic genomic effects, E2 rapidly alters the function of receptors and the activity of second messengers through membrane estrogen receptors such as the typical membrane-associated ERα and ERβ as well as G protein-coupled estrogen receptor 1 (GPER1) (3). E2 promptly regulates glutamatergic neurotransmission and synaptic plasticity. For instance, E2 decreases AMPA miniature excitatory postsynaptic current frequency within minutes (4). Surface trafficking of glutamate receptors such as AMPA receptors (AMPARs) plays critical roles in excitatory neurotransmission (5) and synaptic plasticity (6). However, whether E2 affects AMPAR surface trafficking is unknown. To examine the effect of E2 on the surface trafficking of the glutamate receptor, we applied E2 to live neurons (differentiated from PC12 cells) (Supplementary Fig. 1a, b, c). We imaged single endogenous GluR2-AMPA, which is the most abundant AMPAR subunit in neurons (7), and mGluR1, a metabotropic glutamate receptor involved in rapid membrane action of E2 (8), using ATTO-labeled antibodies and live-cell total internal reflection fluorescence (TIRF) microscopy (Fig. 1a; Supplementary Movies 1-4).

**Figure 1.**
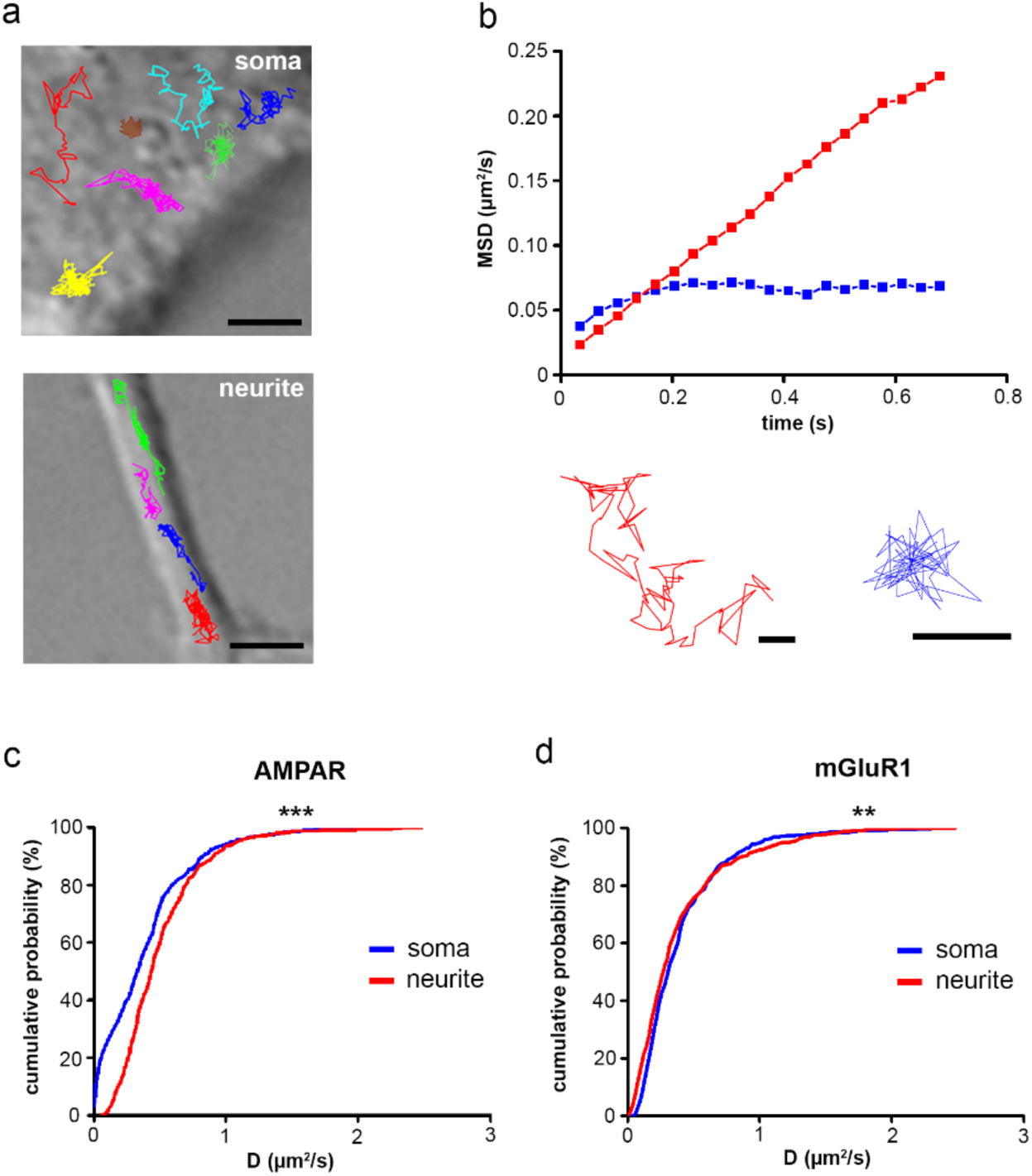
Single-molecule imaging of ATTO 488-labeled GluR2-AMPAR and mGluR1 receptors on neurites and on somas of PC12 neurons. (**a**) Representative trajectories of AMPAR molecules on somas and neurites. Scale bar: 2 µm. (**b**) The mean square displacement functions and trajectories represent AMPAR molecules with Brownian motion (red) and confined motion (blue). Scale bars: 0.1 µm. The cumulative probability functions of D values of AMPAR (**c**) and mGluR1 (**d**) on neurites and on somas are shown (n= 330-500 trajectories). **p<0.01, ***p<0.001

GluR2-AMPAR and mGluR1 molecules exhibited Brownian and confined motion (Fig. 1a, b). The diffusion coefficients of GluR2-AMPAR (D_AMPAR_) and mGluR1 (D_mGluR1_) were significantly greater on neurites than on somas (Fig. 1c, d), indicating that the surface movements of glutamate receptors are faster on neurites.

Application of 100 pM E2 decreased D_AMPAR_ on somas within 5 minutes; however, it did not affect D_AMPAR_ on neurites (Fig. 2a). The effect of E2 was selective for GluR2-AMPAR, since E2 did not change D_mGluR1_ (Fig. 2b; Supplementary Fig. 2b). In contrast, 100 nM E2 decreased D_AMPAR_ on neurites, while D_AMPAR_ on somas remained unchanged (Fig. 2c). Previously, Potier and colleagues demonstrated that 10 nM E2 decreased the diffusion dynamics of NMDA receptor (GluN2B-NMDAR) molecules in hippocampal neurons (9). Our findings indicate that the rapid effect of E2 on the surface trafficking of GluR2-AMPAR is compartment-specific and dose-dependent.

**Figure 2.**
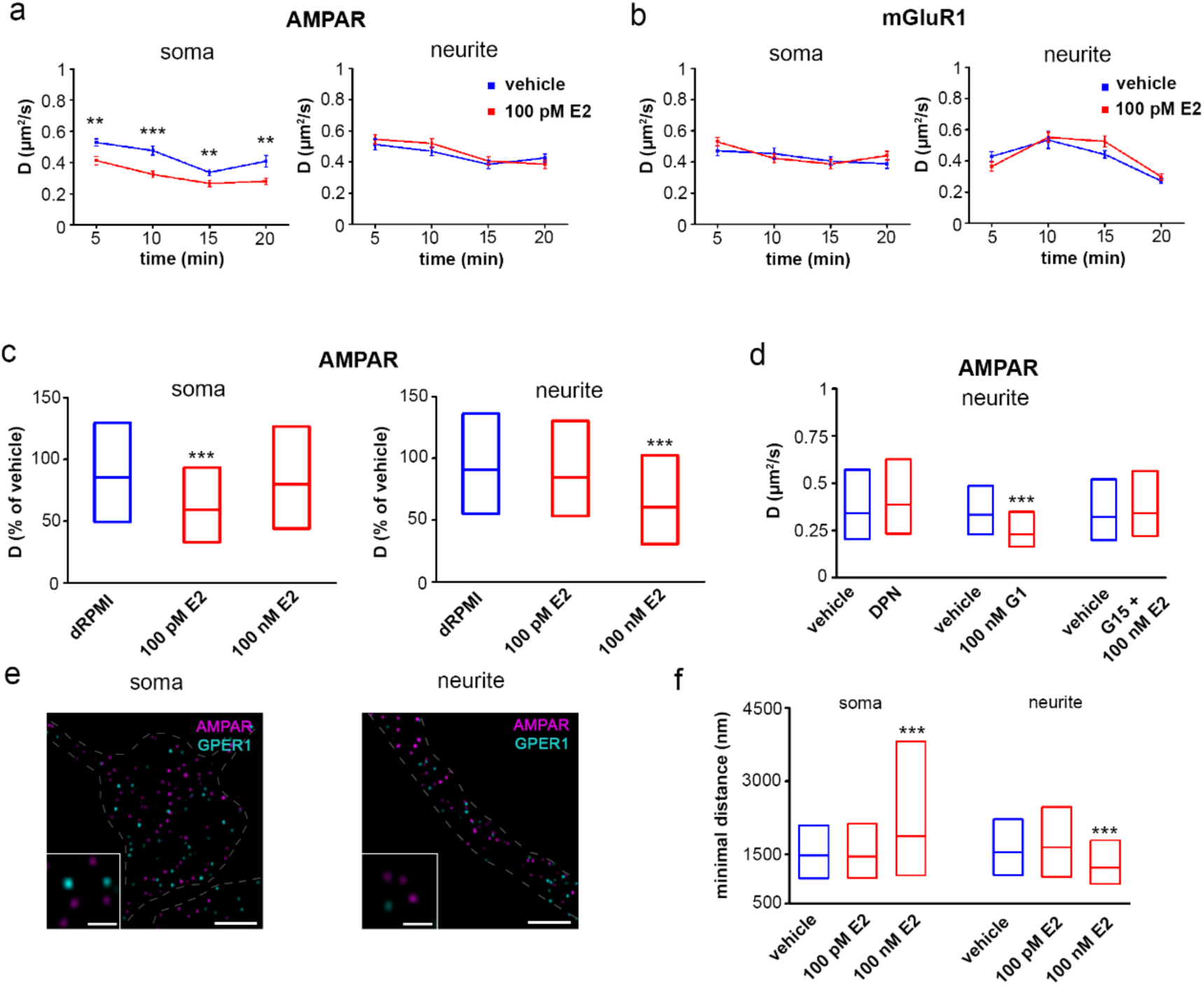
Effect of E2 on the surface trafficking of GluR2-AMPAR and mGluR1. The line graphs depict changes in the diffusion coefficient (D) of GluR2-AMPAR (**a**) and mGluR1 (**b**) molecules after vehicle or 100 pM E2 administration at different time points (mean ± SEM, n=100-128 trajectories per time point). (**c**) Distributions of D (as the percentage of vehicle control) of GluR2-AMPAR (average of 20-minute measurement) in the medium (dRPMI) or after administration of E2 at different concentrations (as the percent of the vehicle treatment expressed as the median ± 25-75%, n= 426-484 trajectories per group). (**d**) Distributions of D values of AMPAR on neurites in the presence of the estrogen receptor β (ERβ) agonist diarylpropionitrile (DPN), a GPER1 agonist (G1) and a GPER1 antagonist (G15) (expressed as the median ± 25-75%, n=427-468 trajectories). (**e**) Stochastic optical reconstruction microscopy images depicting immunolabeled AMPAR (magenta) and GPER1 (cyan) molecules on PC12 neurons. The dashed lines delineate the border between the neurite and soma. (**f**) Distributions of the minimal distance (greater than 500 nm) between GluR2-AMPAR and GPER1 after vehicle or E2 treatment (expressed as the median ± 25-75%, n=203-657). Scale bar: 2 µm; insert scale bar: 0.5 µm. **p<0.01, ***p<0.001

Our PCR results revealed that PC12 neurons express ERβ and GPER1 but not ERα (Supplementary Fig. 2a). Although addition of 10 pM diarylpropionitrile (DPN; an ERβ agonist) (10) had no effect on the surface trafficking of GluR2-AMPAR molecules (Fig. 2d; Supplementary Fig. 2c), 100 nM G1 (a GPER1 agonist) (11) mimicked the effect of 100 pM E2 on D_AMPAR_ in neurites (Fig. 2d) without affecting somatic GluR2-AMPAR (Supplementary Fig. 2c). Prior application of 1 µM G15 (a selective GPER1 antagonist) (11) blocked the effect of E2 on both neurites (Fig. 2d) and somas (Supplementary Fig. 2c). These findings indicate that E2 rapidly decreases the surface trafficking of GluR2-AMPAR molecules via GPER1. In addition, the results of our stochastic optical reconstruction microscopy demonstrated that the median minimal distance between AMPAR and GPER1 molecules (MD_AMPAR-GPER1_) was approximately 1500 nm (Fig. 2e, f; Supplementary Fig. 2d). This result suggests that AMPAR and GPER1 molecules do not form protein complexes in the membrane. To test whether the effect of E2 is accompanied by the redistribution of GPER1 molecules and AMPARs, MD_AMPAR-GPER1_ was calculated after E2 treatment. Although 100 pM E2 did not have an effect, 100 nM E2 increased MD_AMPAR-GPER1_ in somas but decreased MD_AMPAR-GPER1_ in neurites, indicating that AMPARs and GPER1 molecules became closer on neurites than on somas after treatment (Fig. 2; Supplementary Fig. 2d). Our findings suggest that GPER1 with bound ligand can remotely alter the surface trafficking of AMPAR and that E2 has a dose-dependent and compartment-specific effect on MD_AMPAR-GPER1_.

The actin network is essential in the organization of neuronal compartments and plays a crucial role in membrane receptor trafficking (12). We used genetic labeling of cortical actin and structured illumination microscopy super-resolution imaging to reveal the effect of E2 on the cortical actin network. Our results from maximum intensity projections of a series of time-lapse images of cortical actin (see Supplementary Methods) revealed that E2 treatment alone did not change cortical actin dynamics in CHO cells (Fig. 3a and Supplementary Fig. 3a). To determine the role of cortical actin in the effects of E2, we treated cells with the actin polymerization blocker latrunculin A (latA, 1 µM). The cortical actin network dynamics in CHO cells and the morphology of the cortical actin network in PC12 neurons were disrupted by latA administration (Fig. 3a, b; Supplementary Fig. 3a; Supplementary Movies 5, 6). In single-molecule imaging experiments, 10 minutes of latA administration increased D_AMPAR_ on somas without affecting D_AMPAR_ on neurites in PC12 neurons (Fig. 3c). Pretreatment with latA diminished the effect of E2 on the surface trafficking of GluR2-AMPAR molecules on somas (Fig. 3d). These data indicate that cortical actin is a critical factor in the compartment-specific surface trafficking of AMPAR and plays a pivotal role in the E2-induced effects on the surface diffusion of somatic AMPAR. Our results also suggest that cortical actin is important in the remote effects of GPER1 on AMPAR.

**Figure 3.**
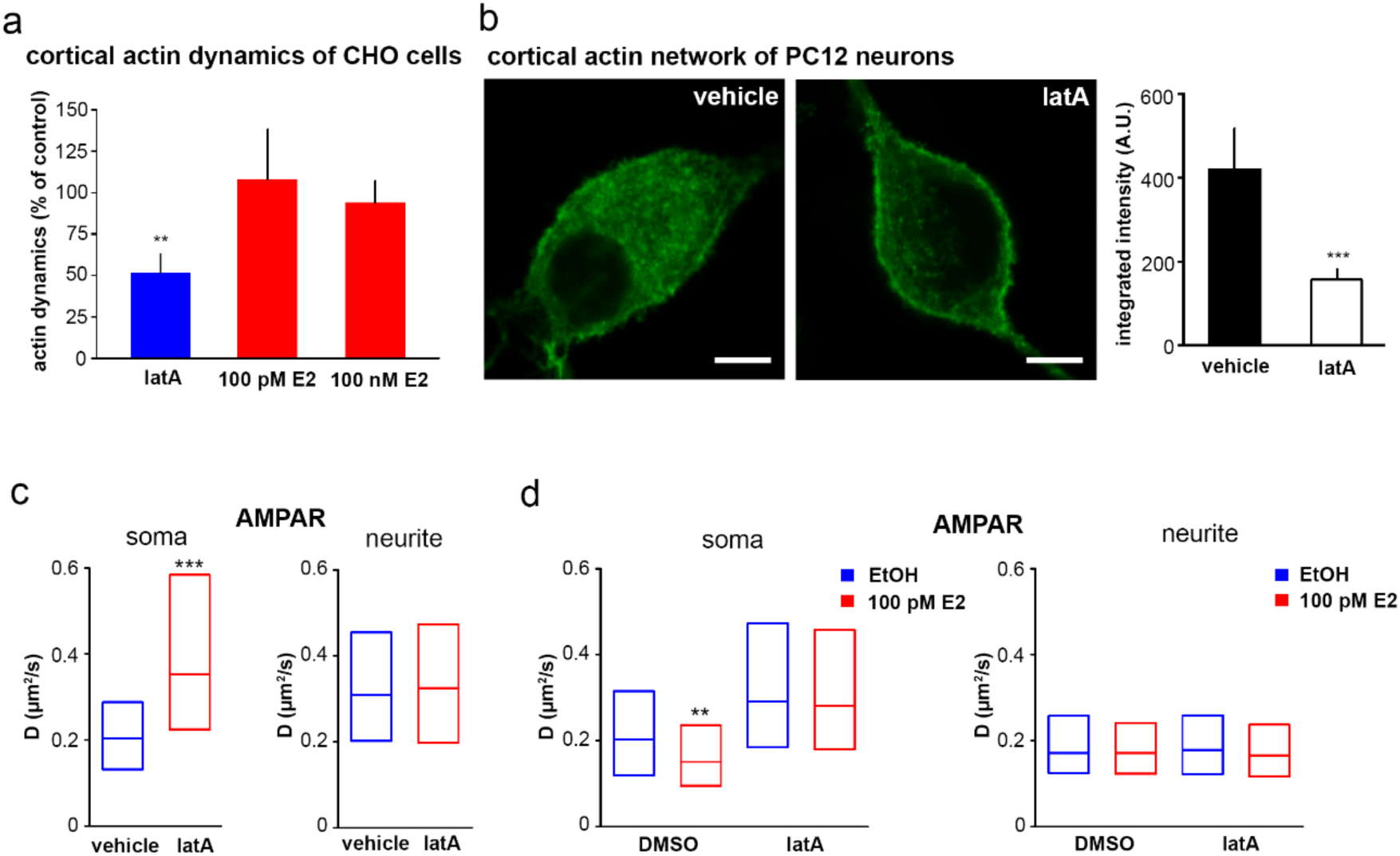
The role of the cortical actin network in the rapid effect of E2. (**a**) The actin polymerization blocker latrunculin A (latA) decreased the dynamics of the cortical actin network in CHO cells (as % of control: 51 ± 11) (for the methods of analysis of cortical actin network dynamics, see Supplementary Methods). Neither 100 nM nor 100 pM E2 affected actin dynamics (as % of control: 100 pM E2 = 108 ± 30; 100 nM E2 = 93 ± 14) (n=5-6 cells per group) (**b**) Confocal images depicting the Alexa Fluor 488 phalloidin-labeled cortical actin network in PC12 neurons after vehicle or 10-minute 1 µM latA treatment in PC12 neurons. The bar graph shows the decreased integrated density of the cortical actin network after latA exposure (right, n=3 cells per group (3 regions of interest per cell)). Scale bar: 5 µm. (**c**). LatA treatment increased the diffusion coefficient (D) of GluR2-AMPA molecules on somas with no effect on the molecules on neurites (expressed as the median ± 25-75%, n = 307-507 trajectories). (**d**) On somas, 100 pM E2 decreased the D values of GluR2-AMPAR molecules. This effect was blocked in the presence of latA. Ineffectiveness of E2 action on neurite was not affected by latA treatment (expressed as the median ± 25-75%, n = 119-205 trajectories). **p<0.01, ***p<0.001

In summary, this study shows the first evidence that E2 decreases the surface movement of GluR2-AMPA in a dose-dependent and compartment-specific manner. E2 alters the surface trafficking of GluR2-AMPA molecules via GPER1, and its action depends on the integrity of the cortical actin network. These observations provide molecular evidence for the remote interaction of GPER1 and AMPAR in neuronal membranes.

## Acknowledgments

This work was supported by the Hungarian Brain Research Program (KTIA_NAP_13-2014-0001,20017-1.2.1-NKP-2017-00002), the Hungarian Scientific Research Fund (OTKA; 112807), and the European Union and was cofinanced by the European Social Fund under the following grants: EFOP-3.6.1.-16-2016-00004 (Comprehensive Development for Implementing Smart Specialization Strategies at the University of Pécs), EFOP 3.6.2-16-2017-00008 (The Role of Neuro-inflammation in Neurodegeneration: From Molecules to Clinics) and ÚNKP-18-3-III (New National Excellence Program of the Ministry of Human Capacities). We thank Dr. Imre Farkas for the generous gift of DPN. The authors wish to thank the Nikon Microscopy Center at the Institute of Experimental Medicine (IEM), Nikon Austria GmbH and Auro-Science Consulting Ltd. for kindly providing microscopy support.

---

**Supplementary Figure 1.**
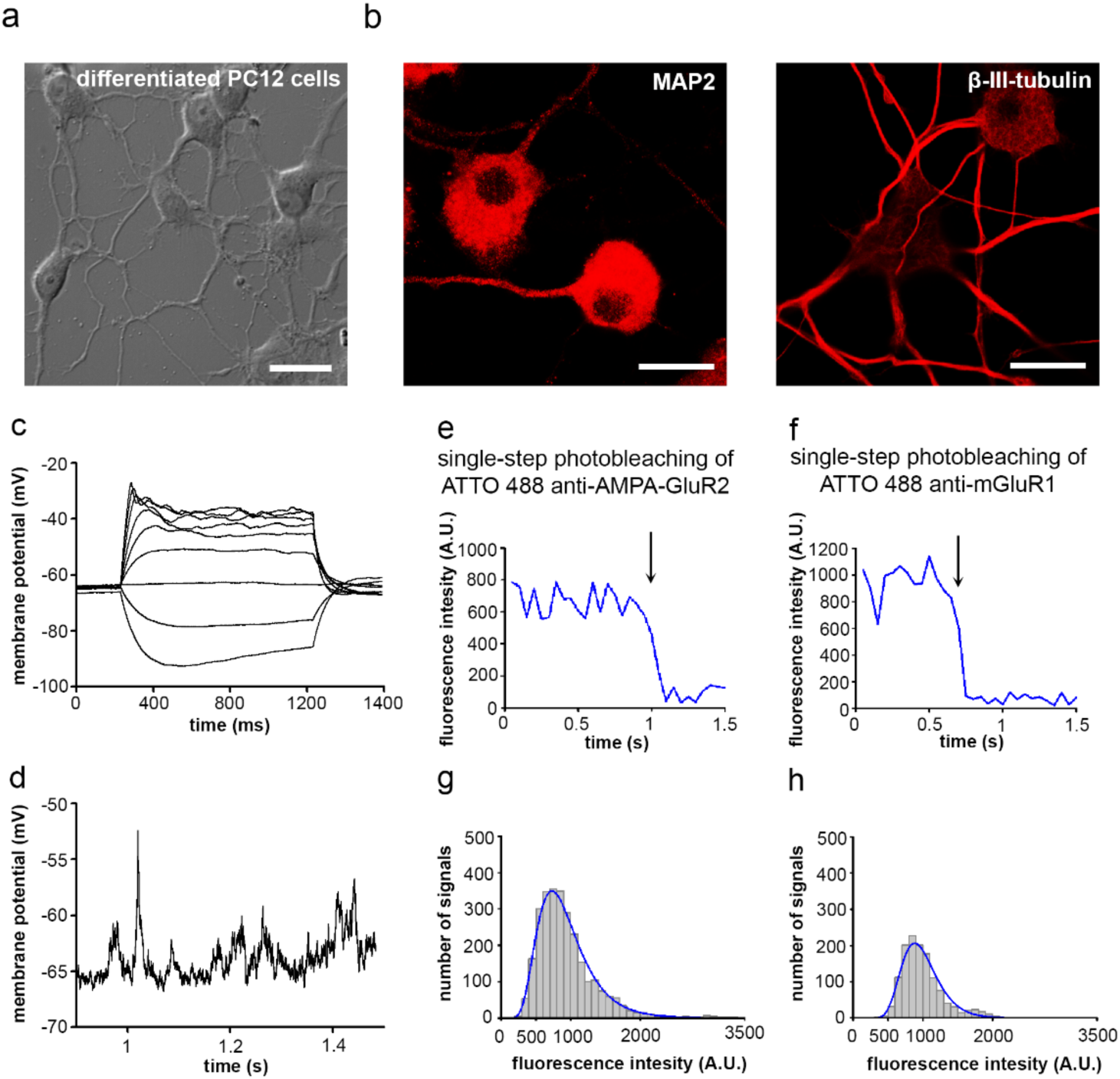
Characterization of differentiated PC12 cells. (**a**) Photomicrograph depicting PC12 cells after 4 days of NGF treatment and (**b**) immunofluorescence staining of microtubule-associated protein 2 (MAP2) and β-III tubulin. Scale bars: 20 µm. (**c, d**) Representative recording of a whole-cell current-clamp step protocol obtained from PC12 neurons. (**c**) Application of increasing current steps in PC12 neuronal culture resulted in action potential generation. (**d**) Trace representing membrane potential changes in PC12 neurons. (**e, f**) Intensity profiles of a single ATTO 488-labeled GluR2-AMPAR and mGluR1 signal. The arrows indicate single-step photobleaching. (**g, h**) Histogram showing the intensity value of every spot found in a recording superimposed with a single fitted Gaussian curve (blue line).

**Supplementary Figure 2.**
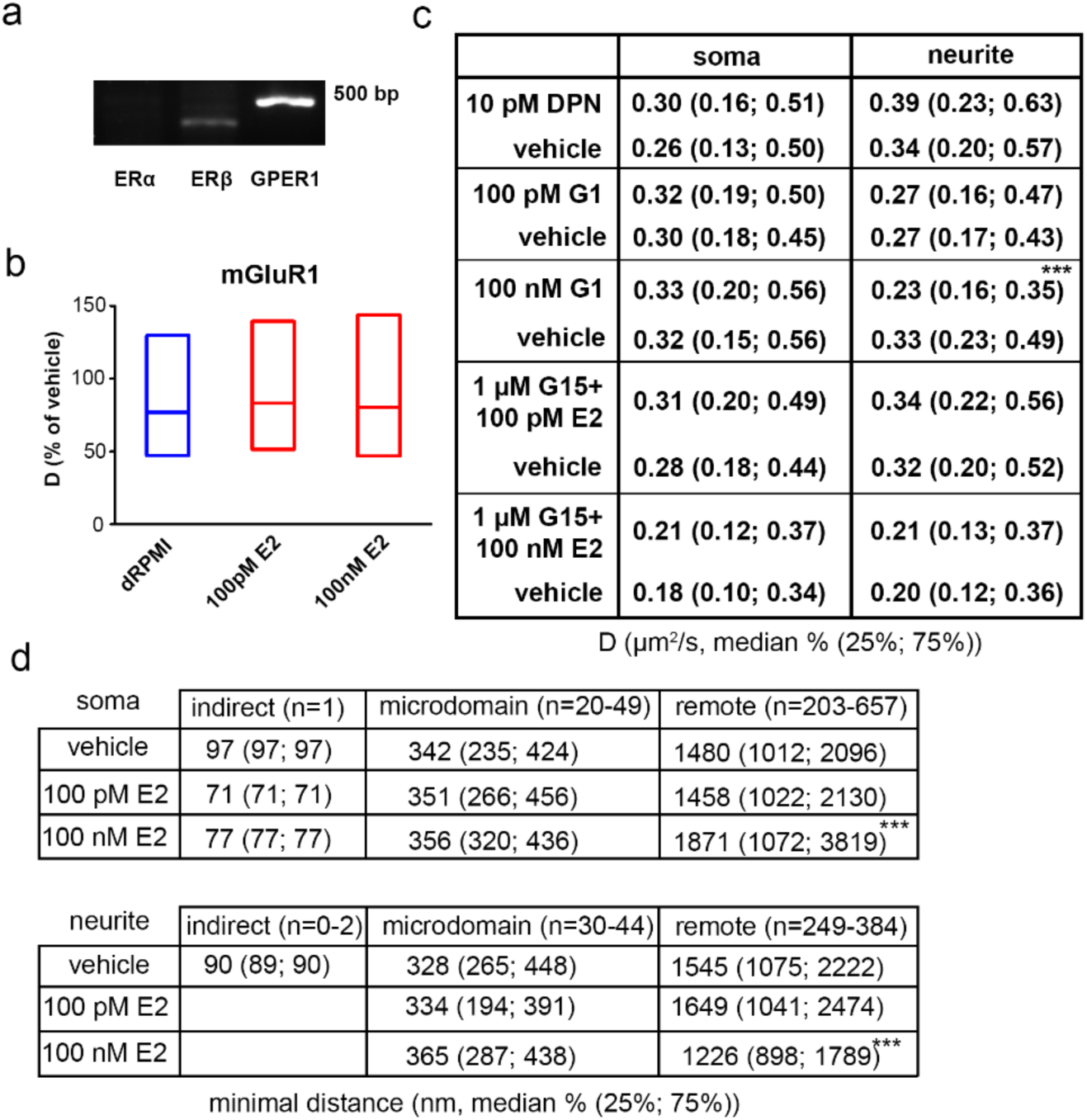
Effect of estrogen receptor modulation on surface trafficking of AMPAR. (**a**) Representative PCR gel electrophoresis image depicting the expression of estrogen receptor mRNA (with a lack of expression of estrogen receptor alpha (ERα), estrogen receptor beta (ERβ), and G protein-coupled estrogen receptor 1 (GPER1)) in PC12 neurons. (**b**) Effect of E2 on the D value of mGluR1 (as the percent of vehicle treatment expressed as the median ± 25-75%, n= 840-860 trajectories per group). (**c**) Table showing the effect of ERβ agonist (DPN), GPER1 agonist (G1) and GPER1 antagonist (G15) treatment on D (µm^2^/s) values of GluR2-AMPAR molecules (expressed as the median ± 25-75% (in brackets), n=411-525 trajectories). (**d**) Table of the distributions of each minimal distance interaction type, including indirect (50-100 nm), microdomain (100-500 nm) and remote (500< nm), between GluR2-AMPAR and GPER1 on PC12 neurons after vehicle or E2 treatment (the minimal distance is expressed as the median ± 25-75% (in brackets)).

**Supplementary Figure 3.**
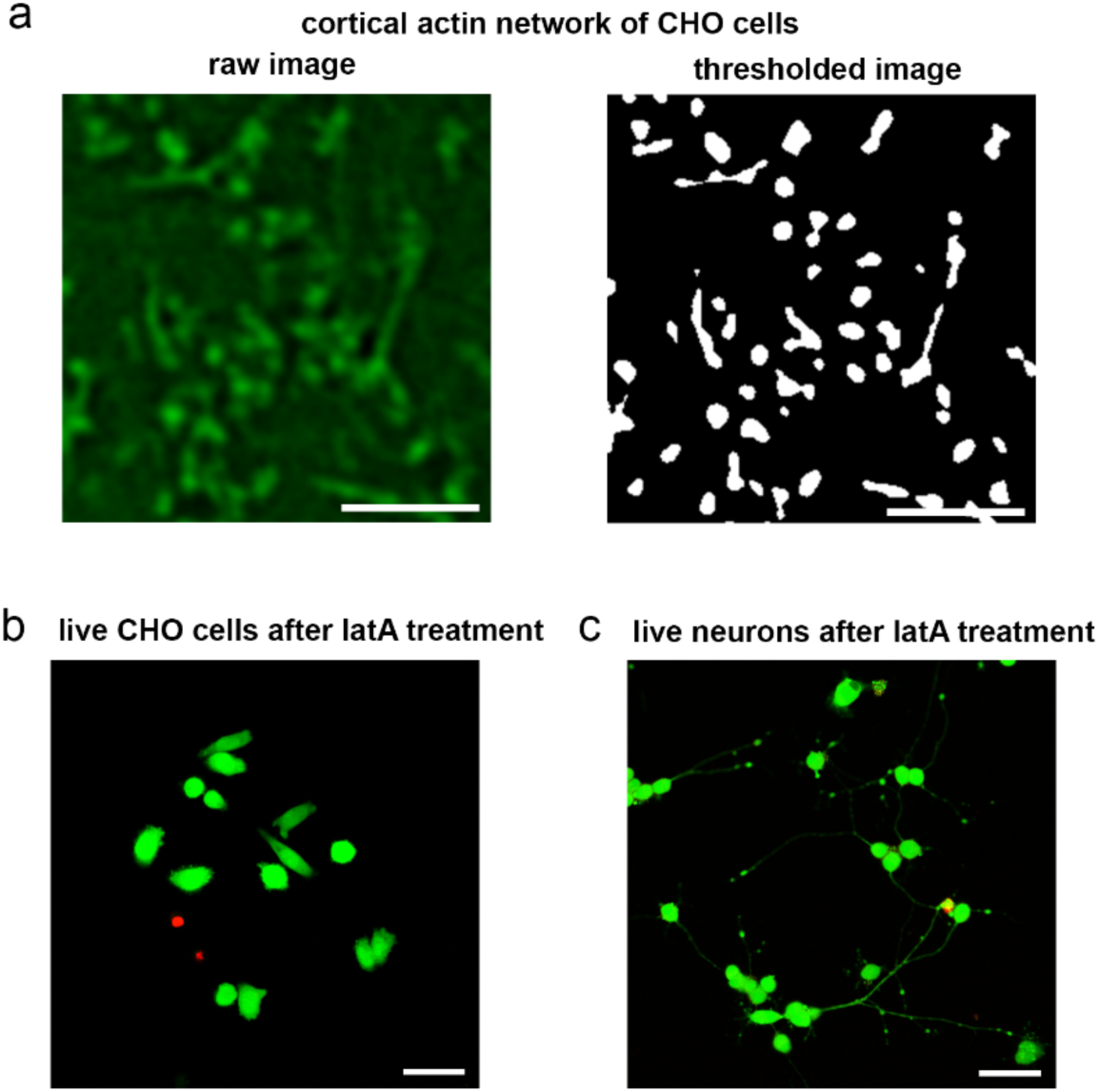
(**a**) Raw (left) and processed (right) immunofluorescence images obtained with structured illumination microscopy depicting the Lifeact-labeled cortical actin network in CHO cells. Scale bar: 2 µm. (**b, c**) The viability of CHO cells and PC12 neurons labeled with a LIVE/DEAD^TM^ Viability/Cytotoxicity Kit (Thermo Fisher Scientific) is shown in fluorescence images 15 minutes after latrunculin A (latA) treatment. The green cells are viable, whereas the red cells have lost their plasma membrane integrity. Scale bar: 50 µm.

## Supplementary Movies

**Movie S1, S2**. Movies of ATTO 488-labeled GluR2-AMPAR on the neurites (S1) and somas (S2) of live PC12 neurons. The recordings were made in TIRF mode with a 34-ms acquisition time and are displayed at 29 fps. Scale bar: 5 µm.

**Movie S3, S4**. Movies of ATTO 488-labeled mGluR1 on the neurites (S3) and somas (S4) of live PC12 neurons. The recordings were made in TIRF mode with a 34-ms acquisition time and are displayed at 29 fps. Scale bar: 5 µm.

**Movie S5, S6**. Movies of the mEmerald-Lifeact-7-labeled cortical actin network before (S5) and after (S6) 1 µM latA treatment. The recordings were made with SIM with a 4-second acquisition time and are displayed at 10 fps. Scale bar: 2 µm.

## Supplementary Methods

### Cell culture and neuronal differentiation

For single-molecule imaging of glutamate receptors, rat pheochromocytoma cells (PC12, Sigma-Aldrich) were differentiated into neurons. PC12 cells were plated at a density of 2×10^3^ cells/cm^2^ on collagen IV-coated 35-mm glass-bottom dishes (MatTek Corporation) in phenol red-free RPMI 1640 medium supplemented with 10% HS, 5% FBS and 2 mM L-glutamine (culture RPMI, cRPMI). Twelve hours after plating, the medium was replaced with phenol red-free RPMI 1640 medium supplemented with 1% HS, 2 mM L-glutamine and 50 ng/ml nerve growth factor (NGF-2.5S, Sigma-Aldrich) (differentiation RPMI, dRPMI). The cells were fed with dRPMI after 2 days and used for imaging after 4 days of differentiation (Supplementary Fig. 1a). CHO cells were cultured in phenol red-free F12 medium supplemented with 10% FBS and 2 mM L-glutamine (culture F12, cF12) at 37°C in an incubator at 95% relative humidity with 5% CO_2_ and plated on glass-bottom dishes (MatTek Corporation) 24 hours before the experiments.

### Validation of neuronal differentiation with immunohistochemistry and electrophysiology

To validate the neuronal differentiation of PC12 cells, MAP2 and β-III tubulin immunocytochemistry were performed. Cells were fixed in 4% paraformaldehyde for 15 minutes and permeabilized with 0.03% Triton X-100 for 30 minutes after 4 days of differentiation. The cells were then incubated with either mouse anti-MAP2 antibody (1:1000, MAB3418, Millipore) or mouse neuron-specific β-III tubulin antibody (1:1000, MAB1195, RD Systems) overnight at 4°C before being incubated with biotinylated donkey anti-mouse F(ab’)_2_ (1:200, Jackson ImmunoResearch) and Alexa Fluor 647-conjugated streptavidin (1:2000, Thermo Fisher Scientific). A confocal laser-scanning microscope (Zeiss LSM710, 100X) (CLSM) was used to detect MAP2 and β-III tubulin immunoreactivity. A helium neon laser with a 633-nm wavelength was used to excite Alexa Fluor 647. The diameter of the pinhole aperture was 80 nm, which resulted in an optical thickness of 2 µm at a wavelength of 690 nm (Supplementary Fig. 1b).

The electrophysiological properties of neurons differentiated from PC12 cells were tested using whole-cell patch-clamp recording. Patch pipettes (1.5 mm outer diameter and 1.1 inner diameter) with a resistance of 6 MΩ were pulled from borosilicate glass capillaries with a micropipette puller (Sutter Instruments). The pipette recording solution contained (in mM) 10 KCl, 130 K-gluconate, 1.8 NaCl, 0.2 EGTA, 10 HEPES, and 2 Na-ATP, and the pH was adjusted to 7.3 with KOH. All recordings were performed at 32°C in a chamber perfused with oxygenated artificial cerebrospinal fluid (ACSF) containing (in mM) 2.5 KCl, 10 glucose, 126 NaCl, 1.25 NaH_2_PO_4_, 2 MgCl_2_, 2 CaCl_2_, and 26 NaHCO_3_. Whole-cell recordings were made with an Axopatch 700B amplifier (Molecular Devices) using an upright microscope (Nikon Eclipse FN1) equipped with infrared differential interference contrast (IR-DIC) optics. Cells with access resistance below 20 MΩ were used for analysis. Signals were low-pass filtered at 5 kHz and digitized at 20 kHz (Digidata 1550B, Molecular Devices). Acquisition and subsequent analysis of the data were performed using Clampex9 and Clampfit software (Axon Instruments). Traces were plotted using Origin8 software (MicroCal Software, Northampton, MA) (Supplementary Fig. 1c).

### Detection of estrogen receptors

Expression of estrogen receptor α (ERα), estrogen receptor β (ERβ) and the membrane estrogen receptor G protein-coupled estrogen receptor 1 (GPER1) was examined in neurons differentiated from PC12 cells. Total RNA was extracted from neurons with a conventional TRIzol (Thermo Fisher Scientific)-based protocol, and cDNA was constructed using a High-Capacity RNA to cDNA Kit (Thermo Fisher Scientific). For PCR, the following primers were used: ERα, 5’-CGTAGCCAGCAACATGTCAA-3’ and 5’-AATGGGCACTTCAGGAGACA-3’; ERβ, 5’-GAGGTGCTAATGGTGGGACT-3’ and 5’-CTGAGCAGATGTTCCATGCC-3’; and GPER1, 5’-TGCACCTTCATGTCCCTCTT-3’ and 5’-AAGGACCACTGCGAAGATCA-3’ (Supplementary Fig. 2a).

### Glutamate receptor labeling in live neurons

To detect AMPA-GluR2 and mGluR1 molecules in the plasma membranes of PC12 neurons, live-cell immunocytochemistry was performed. Before single-molecule imaging, PC12 neurons were incubated in dRPMI with ATTO 488-labeled antibodies directed against the extracellular N-terminal domain of rat GluR2 (1:100, Alomone Labs) or against the extracellular N-terminal domain of rat mGluR1 (1:100, Alomone Labs) at 37°C for 6 minutes. The specificity of the antibodies was tested with control peptides (GluA2_179-193_ peptide and mGluR1_501-516_ peptide, Alomone Labs), and no immunoreactivity was observed.

### Drug application and cell viability detection

The following drugs were applied immediately before imaging in dRPMI for neurons: 17β-estradiol (E2, Sigma-Aldrich, 100 pM in 10^−5^ % EtOH and 100 nM in 10^−3^ % EtOH); G1, a selective GPER1 agonist (Tocris, 100 pM in 10^−3^ % DMSO and 100 nM in 10^−5^ % DMSO); and diarylpropionitrile (DPN), a selective ER-β agonist (Tocris, 10 pM in 2×10^−5^ % DMSO). To block GPER1, neurons were incubated in dRPMI containing G15, a selective GPER1 antagonist (Tocris, 1 µM in 2×10^−3^ % DMSO) for 10 minutes before E2 application and imaging. The actin polymerization inhibitor latrunculin A (latA, Sigma-Aldrich, 1 µM in 0.1% DMSO) was applied for 5 minutes before E2 addition and imaging. CHO cells were treated with latA (1 µM) or E2 (100 pM or 100 nM) in cF12 for 15 minutes.

The viability of the cells was tested after latA treatment with a LIVE/DEAD viability/Cytotoxicity Kit (Thermo Fisher Scientific) (Supplementary Fig. 3b, c) at the end of experiments according to the manufacturer’s instructions. The results demonstrated that cells retained their plasma membrane integrity until the end of the experiments.

### Single-molecule imaging of glutamate receptors using total internal reflection fluorescence microscopy (TIRFM)

Single-molecule imaging of labeled glutamate receptors was carried out on an Olympus IX81 fiber TIRF microscope equipped with Z-drift compensation (ZDC2) stage control, a plan apochromat objective (100X, NA 1.45, Olympus) and a humidified chamber heated to 37°C containing 5% CO_2_. The dish containing the neurons was mounted in the humidified chamber of the TIRF microscope immediately after in vivo labeling. A 491 diode laser (Olympus) was used to excite ATTO 488, and emission was detected above the 510 nm emission wavelength range. The angle of the excitation laser beam was set to reach a 100 nm penetration depth of the evanescent wave. A Hamamatsu 9100-13 electron-multiplying charge-coupled device (EMCCD) camera and Olympus Excellence Pro imaging software were used for image acquisition (Supplementary Movies).

Experiments were performed for 20 minutes. During the measurement period, 20-30 images image series were recorded with 10-second sampling intervals and 33-ms acquisition times. Single-molecule tracking was performed with custom-made software written in C++. The center of each particle was localized by two-dimensional Gaussian fitting. The trajectory for each signal was created by a minimum step size linking algorithm that connected the localized dots in subsequent images. The trajectories were individually checked, and artifacts or tracks shorter than 15 frames were excluded from further analysis. A minimum of 400 trajectories were collected in each experiment from both somas and neurites. To examine the effect of 100 pM E2 or vehicle (EtOH), 100-150 trajectories were collected in every consecutive 5-minute interval up to 20 minutes (0-5; 5-10; 10-15; and 15-20 minutes).

The fluorescence intensity versus time function showed one-step photobleaching representing single ATTO 488 fluorophores. The peak intensity of the intensity spot frequency histogram was similar to that of the step size for photobleaching (Supplementary Fig. 1d, e). These results suggest that most of the spots represented single fluorophores and single receptors.

### Diffusion coefficient (D) calculation with the maximum likelihood method

To calculate D for moving molecules, we first determined the immobile trajectories with MATLAB-based software (TrackArt) (1). Since a zero or negative slope of a line fitted to the first 3 values of the mean square displacement (MSD; µm^2^) curve clearly reveals immobile behavior of an observed trajectory, such line fitting is a commonly used and reliable method to identify and exclude immobile trajectories (2). MSD for each trajectory was calculated with the following equation:

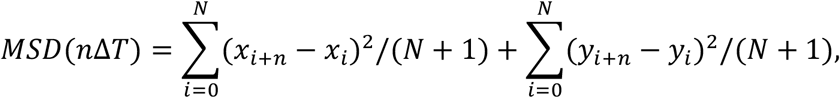

where x_(i)_ and y_(i)_ are the coordinates of the center of the signal, ΔT is the time interval, n is the number of frames within the trajectory, and N is the total number of frames. D (µm^2^/s) for each trajectory was calculated from the slope of the line fitted to the first 3 values of the MSD curve with the following equation:

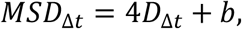

where b is the value at which the function crosses the y axis.

Immobile trajectories (with zero or negative D values) were excluded from further analysis to calculate D values of moving molecules. Based on its asymptotically optimal properties (3), the maximum likelihood method is expected to more precisely estimate D for the moving trajectory of a molecule than other methods. Maximum likelihood estimation (4) was applied to obtain the corresponding diffusion coefficient for each trajectory. Δ_k_ represents the observed displacements 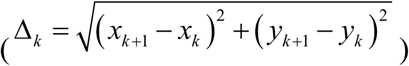 arranged in an *N*-component column vector, where the total number of frames is equal to *N*+1 and *x*_n_ and *y*_n_ are the coordinates of the signal center on the *n*th frame. Σ is the *N* × *N* covariance matrix defined by the following equation:

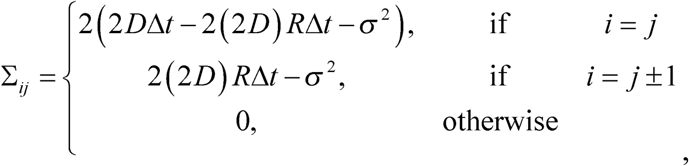

where D is the diffusion coefficient, Δ*t* is frame integration time, *σ* is the static localization noise, and *R* summarizes the motion blur effect; in our case, *R*=1/6 as a consequence of uniform illumination.

The likelihood was defined by the following function:

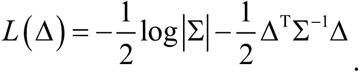

This likelihood function was maximized by a numerical method implemented in MATLAB.

### Colocalization analysis of AMPAR and GPER1 using stochastic optical reconstruction microscopy (STORM)

To examine the probability of interaction between AMPAR and GPER1, STORM imaging was performed. PC12 cells were plated onto poly-D-lysine (PDL)- and laminin-coated coverslips (5) and differentiated into neurons as described above. The neurons were incubated in dRPMI containing vehicle (EtOH) or E2 (100 pM or 100 nM) at 37°C for 10 minutes. Immediately after treatment, AMPA-GluR2 was labeled in live PC12 neurons with anti-AMPA-GluR2 antibody (1:1000, MAB397, raised in mouse, Millipore) at 37°C for 20 minutes followed by fixation in 4% paraformaldehyde. After thoroughly washing, the cells were incubated with anti-GPER1 primary antibody (1:5000, AF5534, raised in goat, Novus Biological) at 4°C for 48 hours. CF-568-labeled donkey anti-goat antibody (1:400, Biotium) was applied at room temperature for 2 hours. Following three consecutive washes, Alexa Fluor 647-labeled anti-mouse antibody was applied at room temperature for 2 hours (1:200, Jackson ImmunoResearch). The coverslips were washed, covered with imaging medium (5) and transferred onto standard glass slides immediately before imaging. Using a CFI Apo TIRF 100X objective, confocal super-resolution images were collected with a Nikon N-STORM C2+ super-resolution system based on the platform of a Nikon Ti-E inverted microscope equipped with a Nikon C2 confocal scan head and an Andor iXon Ultra 897 EMCCD camera. 3D STORM imaging was achieved with the astigmatic imaging method. STORM images were acquired by illuminating the samples with high-powered lasers (561 and 647 nm). Image acquisition and processing were performed using Nikon NIS-Elements AR software with the N-STORM module. The obtained 3D STORM localization points were filtered for the collected photon number, z position (within an axial distance of −300 to 300 nm from the center plane) and local density using VividSTORM software (6). Localization points were selected according to regions of interest (ROIs) manually defined based on the correlated high-resolution confocal images. The clusters of selected localization points were determined by the Density-Based Spatial Clustering of Applications with Noise (DBSCAN) algorithm. The center of mass was calculated for each cluster representing a single molecule. The X, Y and Z coordinates of the center of mass were used to determine the distances between AMPAR and GPER1 molecules (7). The minimal distances between AMPARs and the closest GPER1 molecules were plotted for each treatment (Fig. 2e; Supplementary Fig. 2c). Interactions based on distance were defined as indirect (50-100 nm), microdomain (100-500 nm) (8) and remote (500< nm) (Supplementary Fig. 2c).

### Imaging and analysis of cortical actin network dynamics and morphology using structured illumination microscopy (SIM) and CLSM

Cortical actin dynamics and their response to E2 or latA were examined in live CHO cells transfected (with Lipofectamine 3000, Thermo Fisher Scientific) with a genetically encoded construct specific for filamentous actin, mEmerald-Lifeact-7 (a gift from Michael Davidson, Addgene plasmid #54148). Time-lapse imaging was carried out using a SIM system (Zeiss Elyra S1) equipped with a plan apochromat oil immersion objective (100X, NA 1.46, Zeiss) (Supplementary Movies). The wavelength of the excitation laser was 488 nm, and the emission filter was a 495-550-nm bandpass filter. Time-lapse image sequences of 50 frames were acquired at a frame rate of 0.25 Hz before and after treatment of each cell. Image processing was performed with the Fiji package of ImageJ software (9). The images were bleach corrected, background-subtracted and thresholded. In the thresholded image sequence, the actin signals and the background appeared as white and black pixels, respectively. (Supplementary Fig. 3a). Time-lapse image sequences (50 frames) were defined as image stacks, with each frame representing one slice of the stack. We applied maximal-intensity Z-projection for the image stacks to compress them to a single 2D image. The cortical actin network was characterized quantitatively by the total number of white pixels on the 2D plane (TNWP). To delineate the moving elements of the cortical actin network, static components (mainly stress fibers) were eliminated by subtracting the TNWP of the first 30 images from the TNWP of the whole stack. The pixels were counted in one ROI. They were selected to contain the fewest stress fibers and other relatively static structures possible while covering a large portion (usually more than half) of the cell. ROIs of the same size were used for images before and after treatment.

To validate the effect of latA on neurons, the morphology of the cortical actin network of fixed PC12 neurons was examined after latA administration. After 5 minutes of treatment with 1 µM latA or vehicle (DMSO), PC12 neurons were fixed in 4% paraformaldehyde, permeabilized with 0.1% Triton X-100 for 30 minutes and incubated with Alexa Fluor 488 phalloidin (1:200, Thermo Fisher Scientific) for 45 minutes at room temperature. Imaging was performed with a Zeiss CLSM (see above for parameters). Three ROIs (ROI size: 10.86 µm^2^) on each cell were selected, and the average integrated density was calculated for 3 latA-treated and 3 vehicle-treated PC12 neurons (Fig. 3b).

### Statistics

The D values are expressed as the mean ± SEM (in the time series), the mean ± SEM as a percentage of the vehicle value and as the median ± 25-75% (see figure legends). To compare the distributions of D values, the Kolmogorov-Smirnov test was used. The integrated densities of Alexa Fluor 488 phalloidin were plotted as the mean ± SD and compared with a two-sample t-test. Comparison between values for actin dynamics was performed with a paired t-test. Statistical differences were considered significant at p<0.05. All statistical analyses were performed with Statistica 13.3 for Windows (TIBCO).

